# Cryo-EM of a nucleotide-polymerizing ribozyme enables its predictive improvement

**DOI:** 10.64898/2026.08.14.744467

**Authors:** Deni Szokoli, Jason Hingey, Vivian Wu, Daniel B. Haack, Nicholas Spellmon, Boris Rudolfs, Adamo Mancino, Zhiheng Yu, Navtej Toor, Rhiju Das

**Affiliations:** Department of Biochemistry, Stanford University School of Medicine; Stanford, CA, USA; A-Form Solutions, Inc.; San Diego, CA, USA; Howard Hughes Medical Institute; Stanford, CA, USA; Department of Chemistry and Biochemistry, University of California San Diego; La Jolla, CA, USA; Janelia Research Campus, Howard Hughes Medical Institute; Ashburn, VA, USA

## Abstract

Ribozymes capable of self-replication from nucleotides would have been central to the hypothesized RNA World. The leading laboratory models for such molecules were converted from a class I ligase by *in vitro* evolution but then developed without 3D structures. Here, scaffolded cryo-EM of the substrate-free tC19Z RNA polymerase ribozyme at 3.1 Å resolution shows how this conversion was achieved. An accessory domain evolved from random sequence grips the ancestral ligase through a loop-loop contact, a seam of magnesium ions, and a six-base stack, and rebuilds the ligase’s substrate binding site from different residues of its own. A previously unrecognized pairing, present before substrate binds, sequesters the 5′ end that must otherwise pair with the template. Compensatory mutations to the ribozyme and template, designed to break this ectopic pairing, increase the extension rate. These results suggest that accelerating RNA structure determination may speed progress toward nucleotide-based self-replication.

## Introduction

In the hypothesized RNA World, which is believed to have preceded modern cellular life, ribonucleic acids were chiefly responsible for the roles that are filled today by DNA and proteins (*1–3*). At this time, the replication of genetic information would necessarily have been carried out by a replicase ribozyme, likely through the catalysis of templated ribonucleic acid polymerization (*2*). Many attempts have been made at recreating such a ribozyme capable of self-replication in the laboratory (*4–9*). The most extensively developed are the RNA polymerase ribozymes (RPRs) (*10–18*) that catalyze the extension of RNA primers with nucleotide triphosphate (NTP) monomers, analogous to protein-based replicases. All NTP-utilizing RPRs have been derived by *in vitro* evolution (*18*) from the same ancestral class I ligase (cIL) ribozyme, which was itself the product of *in vitro* selection from a random library (*19, 20*). This conversion from a single-step ligase to a multi-step polymerase was achieved by stripping the ligase of its substrate-binding domains and appending to its 3′-end a random library, from which the NTP-binding “accessory domain” evolved (*18*). Later, the resulting Pol 1 ribozyme would serve as the starting point for the development of the tC19Z RPR, which improved on the sequence generality and primer-template affinity of the ribozyme, allowing it to copy an active hammerhead ribozyme and extend a primer by 95 nucleotides in the same time that the parental Pol 1 could only achieve 8 nucleotide additions (*11*). This improvement required only four point mutations compared to Pol 1 and a short 5′-extension that paired with the template, brought together by rational design guided by secondary structure modeling (*11*). tC19Z would later serve as the starting point for a series of *in vitro* selections which would yield the 24-3, 38-6, 52-2 and most recently the 71-89 RPRs (*12–15*). The latter three are capable of synthesizing active variants of the ancestral ligase, and the 71-89 RPR can replicate and maintain the genetic information of a population of evolving hammerhead ribozymes (*13–15*). Alternative lineages based on assembly of pre-polymerized trinucleotides have been able to make full-length copies of themselves (*9, 21*), but still no RPR can copy itself from single nucleotides.

Crystal structures of the cIL ribozyme established the catalytic core shared by the ligase and its polymerase descendants (*22, 23*), and cryogenic electron microscopy (cryo-EM) of a heterodimeric triplet polymerase ribozyme showed this core to be retained in a ligase descendant with polymerase activity (*24*). However, ribozymes capable of polymerizing nucleotide triphosphates have been refractory to experimental 3D structure determination (*25*). We recently developed a method to enhance cryo-EM of small structured RNAs by fusing them to a highly ordered group II intron scaffold (*26*). Here we determine the structure of tC19Z, the ribozyme from which the modern polymerase lineages descend (*11*), in its substrate-free state at 3.1 Å resolution. This structure reveals long-range RNA-RNA interactions between the catalytic domain (previously called the ligase core domain) and the accessory domain, and accounts for numerous biochemical observations (*27, 28*). It also reveals an ectopic interaction, in which the ribozyme’s template-binding 5′ end is sequestered in a pairing with its own interdomain linker. Mutations designed to release this pairing increase the extension rate, without screening or selection. Substrate-free states may therefore be as informative as catalytically engaged ones for ribozyme improvement, and we conclude by setting out predictions that a substrate-bound structure would test.

### Two-domain structure of the RNA polymerase ribozyme tC19Z

Because tC19Z is relatively small for single-particle cryo-EM (< 200 nt), we embedded the polymerase ribozyme within a circularly permuted group II intron scaffold (*26, 29*). This scaffolding strategy benefits cryo-EM reconstruction by increasing the particle size, reducing orientation bias, and enabling a standardized sample- and grid-preparation workflow. Use of this technique allowed us to solve the tC19Z structure in the apo state at 3.1 Å resolution (**Movie S1, Figure 1A, Figure S1**, and **Table S1**). This structure therefore represents the ribozyme’s ground state, prior to engagement with primer, template, or NTP.

**Figure 1.**
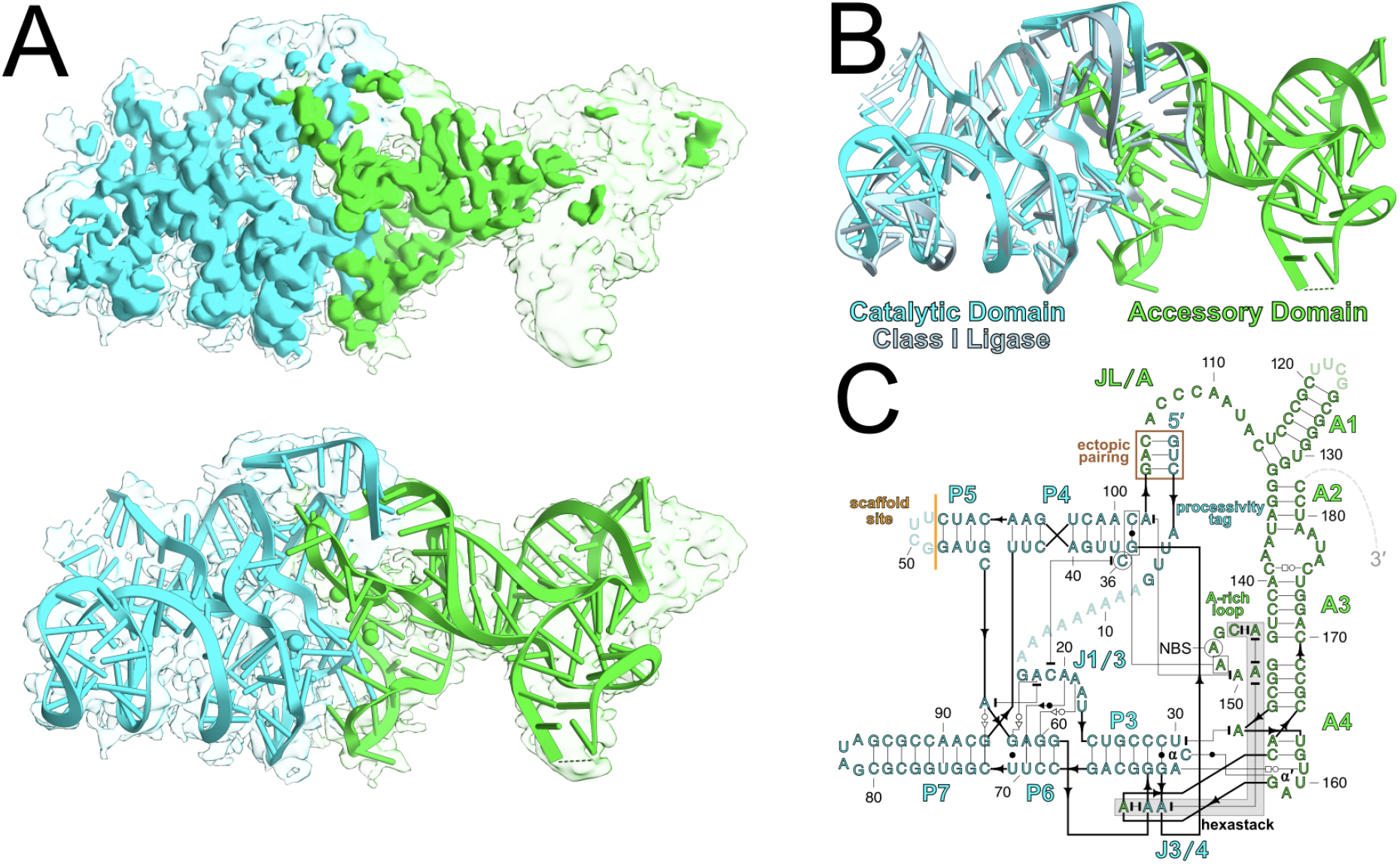
Structure of tC19Z RNA polymerase ribozyme. A) tC19Z map and model. Top: low and high threshold representations of map overlaid. Bottom: low threshold map overlaid with tC19Z model. B) tC19Z model overlaid with corresponding residues in the cIL (PDB 3R1L, light blue) (*23*). C) Secondary structure of tC19Z highlighting both preserved and newly identified tertiary contacts. An ectopic pairing between the processivity tag and JL/A is boxed in brown. A hexastack is highlighted in gray. Dashed lines and grayed-out sequences are unresolved in the cryo-EM maps. The cutoff point where tC19Z was attached to the scaffold is indicated by a solid orange line. Proposed nucleotide binding site is NBS; connectors between nucleotides use Leontis-Westhof annotation (*30*). In (B,C), magnesium ions are shown as spheres. In (A-C), Catalytic domain is shown in cyan. Accessory domain is shown in green.

The overall structure of the catalytic domain has remained unchanged in the tC19Z ribozyme, similar to what has been observed for the triplet polymerase ribozyme evolved from the same cIL ancestor (**Figure S2**) (*24*). The conversion of the ligase to a polymerase has not perturbed the catalytic site, including two catalytic nucleotides that stack on each other, C36 and C19. The surrounding elements that support the formation of the catalytic site (stems P3, P4, P5, P6, and P7, numerous noncanonical pairs, and a Mg^2+^ ion) superimpose precisely between tC19Z and the cIL (**Movie S1, Figure 1B** and **Figure S3**). The structures of some flexible regions could not be solved, such as a run of adenosines between positions 8 and 16 (**Figure 1C**). Overall, the lack of observable change in the 3D fold of the catalytic domain suggests that the catalytic mechanism likely remains unchanged between the polymerase and its ligase ancestor (*23, 25*). While the catalytic domain is largely conserved, the accessory domain represents the major structural addition associated with polymerase function. The accessory domain secondary structure agrees with prior predictions (**Supplementary Text**), but its tertiary fold, evolved from random sequence, forms a high-contact interface with the pre-ordered catalytic domain, similar to RNA aptamers evolved to bind fixed targets, and is described next.

### New tertiary elements span the two-domain interface

The interface between the accessory and catalytic domains includes numerous well-resolved elements mostly not seen in previous structures (**Movie S2** and **Figure S4**). From furthest to closest distance to the ribozyme’s catalytic site (bottom to top in **Figures 1A, 1B**, and **2A**), these elements are: a tertiary loop-loop contact that we term α-α′, a cluster of magnesium ions that we term a ‘magnesium seam’, a collection of six stacked bases that we term a ‘hexastack’, and the proposed NTP binding site.

First, a tertiary contact completes the interface of the accessory and catalytic domains. Chemical mapping suggested that the pentaloop at the tip of A4 might form a long-range contact with the lone-pair triloop in J3/4 (*27*), but its structure remained unknown. This contact, which we name α-α′ (**Figure 2B**), has no structural precedent, based on an automated ARTEM search (*31*) against prior RNA structures (**Methods, Supplementary Text** and **Figure S5A**). All of the loop nucleotides play a part in α-α′ formation (see, e.g., hydrogen bonds as dashed lines in **Figure 2B** and **Supplementary Text**), which may explain why the two loops have remained so well-conserved throughout many generations of *in vitro* evolution (*10–15, 27, 32*). Notably, a C31:G162 Watson-Crick pair connects the triloop and pentaloop of α-α′ (**Figure 2B**), which enabled tests of the functional importance of α-α′ through compensatory mutagenesis experiments. We performed a mutational analysis on the C31:G162 base pair by mutating just one or the other base to induce mispairing and by simultaneously mutating both bases such that the pairing between them would be restored. In polymerization time course assays, disruption of the C:G pair with either a C31G or G162C mutation resulted in reduced activity (**Figure 2C, 2D**). Although the C31G;G162C double mutant did not completely recover wild-type activity, the two mutations were 4.6-fold [95% CI: 3.8–5.6] less deleterious in combination than expected from their individual effects (**Table 1)**. This coupling is the signature expected of two positions that interact directly, and supports the assignment of a C31:G162 pair bridging the catalytic and accessory domains. Incomplete recovery is consistent with additional contacts at this site (**Supplementary Text**)

**Table 1:** Kinetic parameters of tC19Z mutants with fitting-based 95% confidence intervals (CI).

| **Ribozyme** | **P<sub>∞</sub> (%)**<br>**[95% CI]** | **k<sub>obs</sub> (×10<sup>-2</sup> min<sup>-1</sup>)**<br>**[95% CI]** | **k<sub>obs</sub> (rel. to wild type)**<br>**[95% CI]** |
| --- | --- | --- | --- |
| wild type | 85.6<br>[83.9–87.2] | 1.07<br>[1.00–1.14] | 1 |
| ***α-α'* tertiary contact** |  |  |  |
| C31G | 82.1<br>[78.8–85.4] | 0.59<br>[0.52–0.66] | 0.547<br>[0.477–0.626] |
| G162C | 79.3<br>[75.5–83.4] | 0.114<br>[0.101–0.128] | 0.106<br>[0.093–0.121] |
| C31G; G162C | 82.2<br>[80.2–84.2] | 0.287<br>[0.269–0.305] | 0.267<br>[0.234–0.293] |
| C31G × G162C,<br>expected if independent | – | 0.062<br>[0.051–0.077] | 0.058<br>[0.048–0.070] |
| C31G; G162C,<br>cooperativity factor | – | – | 4.6<br>[3.8–5.6] |
| **Release of ectopic pairing** |  |  |  |
| U2A; template-WT | 83.6<br>[81.6–85.7] | 0.251<br>[0.235–0.267] | 0.235<br>[0.214–0.257] |
| WT; template-A8U | n.d. | < 0.054<sup>a</sup> | < 0.051<sup>a</sup> |
| U2A; template-A8U | 93.1<br>[91.2–95.1] | 1.67<br>[1.56–1.80] | 1.56<br>[1.42–1.72] |
| U2A × template-A8U,<br>expected if independent | – | < 0.013<sup>a</sup> | < 0.012<sup>a</sup> |
| U2A; template-A8U,<br>cooperativity factor | – | – | >130<sup>a</sup> |

<sup>a</sup> Bound with 95% confidence; see **Methods**.

**Figure 2.**
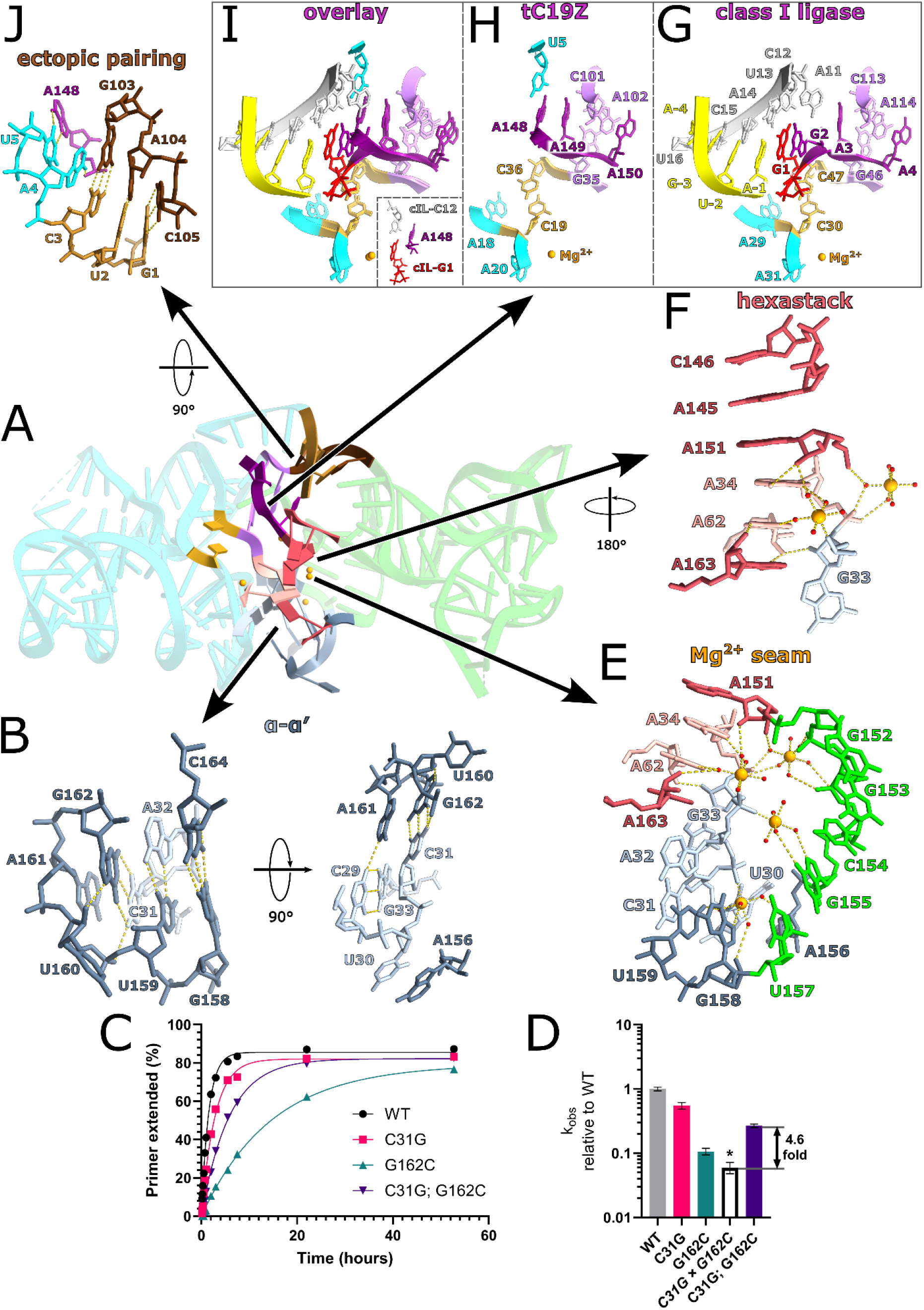
Extensive tertiary features connect the accessory and catalytic domains of tC19Z. A) Global view of the tC19Z RNA polymerase ribozyme. Cyan and green are general residues belonging to the catalytic and accessory domains, respectively. The conserved catalytic cytidine residues are shown in gold. Gray, pink, purple, and brown indicate residues involved in α-α′, the hexastack, the NTP-binding site, and the ectopic pairing, respectively. Within each of these four features, residues from the catalytic domain are a darker shade (e.g., dark gray) while residues from the accessory domain are a lighter shade (e.g., light gray), Mg^2+^ ions are shown in orange, and yellow dashed lines represent hydrogen bonds or metal ion coordination, where relevant. B) Close-up representations of the loop-loop interaction α-α′. The first view focuses on the pentaloop residues from the accessory domain (left), whereas the second view shows all of the triloop residues from the catalytic domain (right). C) Time courses of first nucleotide addition showing the activity of the different mutants that disrupt and restore α-α′ interaction. D) Relative reaction rates of first nucleotide addition normalized to the rate of the wild type ribozyme. White bar (*C31G × G162C*, marked with asterisk) represents the expected rate of the double mutant if single mutants acted independently; error bars show 95% confidence intervals. E) The magnesium seam between the catalytic and accessory domains forms a network of interdomain contacts, with involvement of the hexastack and the α-α′ interaction. F) Close-up view of the hexastack, with participating Mg^2+^ ions shown. G-I) Comparison of (G) cIL (PDB 3R1L) active site and (H) proposed NTP binding site of tC19Z polymerase ribozyme, with (I) overlay showing the mimicry of residues cIL-A3 and cIL-A4 by tC19Z accessory domain residues A149 and A150. The same color scheme used for tC19Z was applied to the corresponding residues in the cIL for ease of comparison. The cIL also has a template shown in silver, primer in yellow, and 5′ nucleotide G1 (analogous to an incoming NTP in the polymerase) in red. The inset in the bottom corner of the (I) shows the position of A148 from tC19Z relative to the position of the NTP-template base pair G1:C12 from the cIL. J) Close-up representation of the ectopic pairing. Immediately adjacent to the three Watson-Crick base pairs of this interaction, A4 also contacts the phosphate of A148 and U5 engages the Hoogsteen edge of A148. See **Figure S4** for cryo-EM map overlays.

Progressing closer to the catalytic site, the α-α′ interaction connects to a cluster of hydrated magnesium ions, which we term a magnesium ‘seam’. This seam bridges the accessory domain and the conserved catalytic domain. This feature is formed by a group of four hydrated Mg^2+^ ions all within 15 Å of one another and positioned along the domain interface, rather than as a compact metal ion core (**Figure 2E, Movie S3**). Three of the ions engage non-bridging phosphate oxygens directly, while all four coordinate water molecules that hydrogen bond to nearby phosphates, ribose hydroxyls, or nucleobase functional groups. Nucleotides C31, A32, G33, A34, and A62 from the catalytic domain and A151, G153, C154, G155, U157, G158, U159, and A163 from the accessory domain all interact with these Mg^2+^ ions or their coordinated water molecules (**Figure 2E, Movie S3**). The seam thus behaves like structural mortar: direct coordination of phosphates by the Mg^2+^ ions provides discrete anchor points, while the ordered waters extend this stabilization across the broader surface of surrounding nucleotides. The result is an extensive solvent-mediated linkage between the accessory and catalytic domains that stabilizes their relative orientation and helps integrate the polymerase-specific accessory addition into the pre-existing ligase fold. The role of four Mg^2+^ ions in mediating this interface may partially explain the high concentration of Mg^2+^ ions (100 mM or higher) required for these ribozymes to function optimally (*10–15, 17, 32, 33*) as well as observations of mutations around this seam in studies that selected for RPR mutants with activity at lower Mg^2+^ concentrations (*34*). In addition to connecting the accessory and catalytic domains of the ribozyme, the magnesium seam also bridges across two tertiary sub-elements of the interface: the α-α′ tertiary contact, described above, and a hexastack, discussed next.

The interface between the ribozyme’s catalytic and accessory domains includes a group of six stacked nucleobases drawn from 5 non-contiguous regions from across the ribozyme, which we refer to as the hexastack (**Figure 2F**). In the order in which they stack, these are C146, A145, A151, A34, A62, and A163; the four accessory domain nucleotides (C146, A145, A151, and A163) flank two adenosines that are holdovers from the ancestral ligase (A34 and A62). Four nucleotides A34, A62, A151, and A163 from this hexastack are also involved in the magnesium seam. While such extensive sets of stacked bases have been observed in prior RNA structures, the stacks typically form from contiguous bases, e.g., in one strand of an A-form helix. An automated scan with ARTEM returned only three arrangements drawn from five separate regions within 2.0 Å RMSD of the hexastack, each failing to reproduce one of its stacking interactions (**Supplementary Text** and **Figure S5B**).

### The NTP binding site recreates the ancestral ligase’s substrate contacts

The structure of the ancestral cIL ligase showed substrate binding to be reminiscent of the active site of proteinaceous polymerases, which bind a primer-template duplex and an incoming NTP that base pairs at the +1 position of the template (*22, 35*). This single Watson-Crick base pair with the template is not sufficient to effectively bind and position the NTP in the active site of proteinaceous polymerases or the cIL ribozyme. Instead, both kinds of enzymes form more extensive binding pockets that support the NTP in base-pairing to the +1 nucleotide of the template and position it for in-line attack of the α-phosphate by the 3′-OH of the primer. In the cIL, the equivalent of the incoming NTP is the triphosphorylated cIL-G1 residue, while its binding pocket consists of the immediately contiguous nucleotides cIL-G2, cIL-A3, cIL-A4, and cIL-A11 (**Figure 2G** and **Figure S3**) (*22*).

In the cIL-derived selection library that would yield the polymerase ribozymes, the P1 stem, the preceding cIL-G2, cIL-A3 and cIL-A4 residues, and the 5′-half of the P2 stem were removed from the library, in a bid to select for a ribozyme that would not need to sequence-specifically bind to the template (*18*). With this, the authors also removed the NTP-binding pocket and the interactions stabilizing the active site. Therefore, any ribozyme selected from this random library would have needed to evolve a new substrate-binding site that could accommodate free NTPs. tC19Z shows a collection of residues near the catalytic C nucleotides that we propose is the ribozyme’s NTP binding site. Adjacent to the hexastack, an accessory domain loop is positioned to bind the NTP substrate (“A-rich loop” in **Figure 1C**). The two adenosines from the ancestral ligase’s substrate binding site (cIL-A3 and cIL-A4), which were excised in the selection library, are structurally replaced in tC19Z by A149 and A150, and are thus contributed by the accessory domain rather than by the catalytic domain (**Figure 2H**). Despite arriving from elsewhere in the molecule, A149 and A150 reproduce three features of the site they replaced: an A-minor interaction with a conserved pair, a splayed backbone conformation, and a stacking interaction with another conserved adenosine (**Figure 2H-I, Supplementary Text**). In an automated ARTEM search, previously observed nucleotide arrangements matched this site geometrically, but the class I ligase itself achieves the closest match at 0.56 Å, compared to a next best match to a generic A-minor interaction of 1.08 Å (**Supplementary Text, Figure S5C**).

The residue immediately preceding these two, A148, is a known determinant of NTP binding (*28*), and its backbone superimposes on the corresponding ligase residue (cIL-G2). Its nucleobase however adopts a different orientation in our substrate-free structure. Instead of an interaction as in the ligase (cIL-A11), the A148 is engaged by U5 in an ectopic pairing (**Figure 2J**), described in the next section. A148 nonetheless remains positioned to contact the base pair formed between an incoming NTP and its templating partner and, with minor structural rearrangements, could stack on the base pair (**Figure 2I inset**).

### Releasing an ectopic pairing increases ribozyme extension rate

Because our structure captures tC19Z before substrate engagement, it also reports on interactions that form only in the enzyme’s ground state. One such interaction involves the single-stranded linker between the catalytic and accessory domains, known as “JL/A” (**Figure 1C**), which is presumed to play a functional role in RNA polymerization. Our structure shows this linker in an unproductive conformation, ectopically base paired with the first three nucleotides of tC19Z, which are a part of the processivity tag (**Figure 1C, Figure 2J**). This 5′ processivity tag arose during *in vitro* evolution and was recognized to base pair to the template to prevent dissociation during primer extension (*11*); however, a subsequent minimization unintentionally brought together a 5′-G with a UC tag dinucleotide to yield a GUC triplet complementary to a GAC trinucleotide in the ribozyme’s own JL/A linker. Considering also that later, more advanced iterations of this ribozyme lineage evolved mutations in the proximal JL/A region that disrupted its ability to pair with the processivity tag (*12*), we suspected that this ectopic base-pairing in tC19Z was competing with the ribozyme’s ability to bind to the template (model shown in **Figures 3A-B, Movie S4**; see **Methods**).

**Figure 3.**
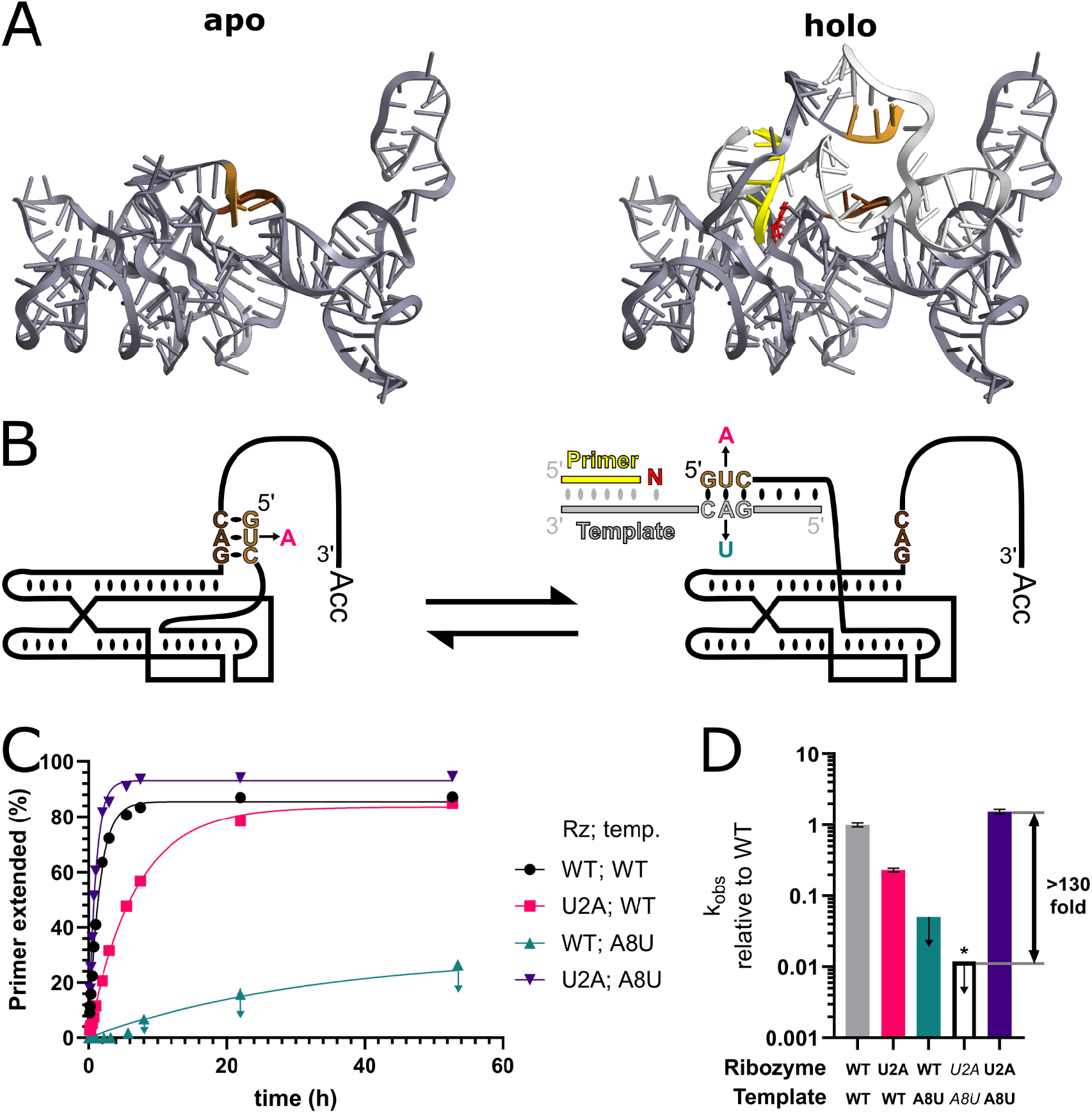
The ectopic pairing between the processivity tag and JL/A. A) Comparison of the tC19Z apo structure (left) and predicted holo structure (right), modeled in Rosetta (**Methods**). Ribozyme residues are shown in dark gray, ectopic pairing in shades of brown as in previous figures, template in silver, primer in yellow, and an incoming GTP in red. B) Schematic of the WT ribozyme and mutants converting between the ectopic and catalytically competent fold. C) Time courses of first nucleotide addition for mutants probing the ectopic base pairing between the processivity tag and JL/A. D) Relative reaction rates of first nucleotide addition normalized to the rate of the wild type ribozyme and template; error bars show 95% confidence intervals. The white bar (*) shows the value expected for the double mutant assuming that the single mutations act independently. Downward arrows in (C,D) on measurements for WT ribozyme with the A8U template signify that they are upper bounds (**Methods**).

Because the ectopic interaction involves Watson-Crick pairings, we were able to rationally design mutations to disrupt it, with the prediction that a designed variant would have increased activity compared to the starting tC19Z sequence. To avoid bias from prior experiments, we made mutations distinct from those seen in later ribozyme lineages, and also chose to make two mutations that we predicted would individually disrupt activity but together would enhance activity (**Figure 3B**). We first tested whether single mutations to either template or ribozyme disrupted RNA polymerization activity (**Figure 3B**). The U2A mutant abolished the ectopic pairing with the JL/A region, but also introduced a single A:A mismatch in its 7-bp interaction with the WT template strand (**Figure 3B**), leading to an overall 4-fold reduction in activity compared to the WT ribozyme (**Figure 3C, 3D**; **Table 1**). When the WT ribozyme was challenged to copy a mutated A8U template, which introduced a single U:U mismatch at the same base pair between template and ribozyme while leaving the ribozyme able to form the ectopic pairing (**Figure 3B**), activity was reduced even more strongly, by at least 19-fold (95% confidence; **Figure 3C, 3D**; **Table 1**). Finally, when the ectopic pairing was disrupted and the pairing of the processivity tag to the template was restored through the combination of both U2A mutation to the ribozyme and A8U mutation of the template (**Figure 3B; Table 1**), the two mutations showed a coupling of greater than 130-fold (with 95% confidence), as expected for mutations that restore base pairing to one another (**Figure 3C, 3D; Table 1** and **Methods**). Moreover, the observed extension rate exceeded that of the WT ribozyme-template combination (1.6-fold; **Figure 3C, 3D**; **Table 1**), reflecting not only restoration of the template-ribozyme base pair but also disruption of the ectopic pairing that competes with template binding. Overall, each of the three predicted outcomes was validated experimentally, establishing the structure supported prediction of mutational effects rather than *post hoc* rationalization.

### Implications for the *holo* state and other RNA polymerase ribozymes

A structure of a single state of a biomolecule can constrain what other states can look like. We therefore set out four predictions that structures of substrate-bound tC19Z, or of other nucleotide-polymerizing ribozymes in this lineage, would test directly. First, the three interdomain elements described above (the α-α′ contact, the magnesium seam, and the hexastack) are expected to be present in substrate-bound states and in descendant polymerase ribozymes (**Figure 2**). The residues involved are conserved through in vitro selections across the ribozyme lineage (*10–17*), and the accessory domain must remain docked for the NTP site to be held against the catalytic center. Rearrangement or loss of any of the three in a substrate-bound structure would falsify this prediction. Second, the mimicry of the ancestral ligase persists on substrate binding. A149 and A150 retain the interaction with G35:C101, the splayed backbone conformation, and the stacking of A150 on A102. The class I ligase structure to which these residues correspond is itself a substrate-engaged state (*22*), so we expect the resemblance to be maintained once substrates are bound. Third, A148 is predicted to contact the base pair formed between the incoming NTP and the templating nucleotide. In our substrate-free structure, the A148 backbone superimposes on the class I ligase’s cIL-G2 while its nucleobase adopts a distinct orientation (**Figure 2I**); although the orientation of A148 in the substrate-bound tC19Z is therefore unclear, with small shifts in the base location, it can stack on the incoming NTP base pair. Fourth, the ectopic pairing between the processivity tag and JL/A should be absent in substrate-bound tC19Z and absent in descendants carrying JL/A mutations (*12*). In these ribozymes the JL/A linker is expected to adopt an alternative conformation, and the 5′ tag should be engaged with template in the *holo* complex.

### Negative design to improve RNA polymerase ribozymes

In this work, we have determined the structure of the tC19Z RNA polymerase ribozyme. The accessory domain of the ribozyme, which evolved to convert the ancestral ligase into a polymerase, forms an aptamer-like grip on the catalytic domain through an α-α′ tertiary contact, a magnesium seam, and a hexastack, all positioning an A-rich loop to mimic the substrate-binding site of the ancestral ligase but with distinct nucleotides. Our *apo* structure also revealed an interaction incompatible with a catalytically competent state, with the ribozyme’s template-binding 5′ end drawn into an ectopic pairing with the ribozyme’s own inter-domain linker. Predictive design of mutations to disrupt this ectopic pairing while restoring template binding led to an increase in extension rates, demonstrating that 3D-structure-guided design of these ribozymes is feasible, in this case based on 3D structure of the substrate-free RNA. Such “negative design” to destabilize ground states or competing states is common in proteins (*36, 37*) and has become increasingly possible for RNA with improvements in cryo-EM (*38*). Beyond the substrate-free ground state, all RNA polymerase ribozymes currently stall or make errors when copying particular sequences and structures. Cryo-EM characterization of these states may enable predictive design of ribozymes that bypass or stabilize them and thereby accelerate development of self-replication from single nucleotides.

## Materials and Methods

### Cryo-electron microscopy

#### Construct design

A group II intron scaffolding strategy was applied to solve the tC19Z structure by cryo-EM. Specifically, a UUCG tetraloop at the end of the tC19Z P5 stem was replaced with the scaffold sequence (**Figure 1C**), thereby increasing the overall size of the particle while preserving the polymerase. The insertion site was chosen based on prior studies showing that L5 loop can be changed without disrupting activity (*22*), as well as to position the scaffold peripherally, thus avoiding disruption of the polymerase active site and native fold. See **Table S2** for sequences.

#### In vitro RNA transcription and purification

The scaffolded tC19Z RNA was transcribed as previously described (*26*), with one modification to the purification, as described in the following. Briefly, the plasmid encoding the fusion construct was linearized at an engineered BamHI site (NEB), and 50 μg of the linearized template was added to a 1 mL transcription reaction (50 mM Tris-HCl pH 7.5, 25 mM MgCl_2_, 5 mM DTT, 2 mM spermidine, 0.05% Triton X-100, and 5 mM of each NTP) containing T7 RNA polymerase and thermostable inorganic pyrophosphatase. The reaction was incubated at 37 °C for 3 hours, after which the DNA template was digested by the addition of Turbo DNase and CaCl_2_ (1.2 mM final) for 1 hour at 37 °C. Proteinase K was then added for an additional hour at 37 °C to degrade the T7 polymerase and DNase, and the resulting solution was centrifuged to remove precipitate. In place of the iterative diafiltration step used previously, the filtered reaction was applied to a HiLoad 16/600 Superdex 200 pg column (Cytiva) on an ÄKTA Pure FPLC equilibrated in running buffer (5 mM Na-cacodylate pH 6.5 and 4 mM MgCl_2_), in order to separate the transcript from residual transcription components. Fractions containing the purified RNA were pooled and supplemented with concentrated MgCl_2_ and KCl stocks to give final concentrations of 10 mM MgCl_2_, 100 mM KCl, and 5 mM Na-cacodylate pH 6.5. The RNA was folded by incubation at 50 °C for 30 minutes followed by 30 minutes at room temperature, and then concentrated to ∼4 mg/mL using a 100 kDa molecular weight cut-off filter..

#### EM sample preparation

3.5 μL of the concentrated RNA was applied to a glow discharged (40 mBar, 15 mA for 45 s using a PELCO easiGlow) Quantifoil R2/1 200-mesh gold grid. The grid was immediately blotted by hand with filter paper (Whatman No.1) before plunge freezing into a 63:37 propane/ethane mixture using a manual plunger.

#### Cryo electron microscopy collection and processing

Cryo-EM data was collected at the Janelia Research Campus on Krios4 equipped with a cold FEG, Selectris-X energy filter and Falcon 4i camera (Thermo Fisher) using SerialEM (Nexperion) with beam-image shift multi-shot acquisition. Movies were recorded in EER format at a physical pixel size of 0.7336Å over a total dose of 50 e^−^/Å^2^.

Data processing was performed in cryoSPARC v4.5. Movies were processed with patch motion correction and contrast transfer function estimation using cryoSPARC’s default settings. Particles were picked using blob picker; after curation, approximately 1 million particles were retained. The initial particle stack was sorted by 2D classification, and 2D projections containing the RNA scaffold were selected for ab initio into 5 classes followed by heterogeneous refinement. Reconstructions containing well-folded tC19Z were selected for further refinement.

To focus particle alignment on the tC19Z insert, the RNA scaffold density was subtracted from particle images using a mask enveloping the scaffold. The remaining sub-particle density of tC19Z was aligned using local refinement to generate a final reconstruction from 150,593 particles. Data processing workflow is illustrated in **Figure S1**. Additional data collection and processing details are listed in **Table S1**. The atomic model of tC19Z was built *de novo* in Coot (*39*), and refined in Coot and PHENIX (*40*). Residues were renumbered to match tC19Z numbering for this manuscript; the PDB deposition uses numbering appropriate for the full length sequence, including the inserted group II intron scaffold. Residues 1-47 and 52-182 of tC19Z correspond to residues 1-47 and 451-581, respectively, in the PDB deposition.

### Ribozyme activity measurements

#### Synthesis and purification of RNA

Transcription templates for ribozyme experiments were ordered as dried gBlocks (IDT Technologies, Coralville, IA) and resuspended to concentrations of 5 ng/μL in IDTE buffer; sequences are provided in **Table S2**. To make dsDNA, 1 μL of each gBlock construct was mixed with 25 μL of NEBNext Ultra II Q5 Master Mix (NEB, M0544S), 2.5 μL of 10 μM phi25_a_tC19Z-FIX-fwd, 2.5 μL of 10 μM tC19Z-rev_methyl, and 19 μL of water. For U2A, a different forward primer, phi25_a_tC19Z-FIX-fwd_U2A, was used. The thermocycler setting was 98 °C for 30 seconds, 30 cycles of 98 °C for 10 seconds, 62 °C for 30 seconds, and 72 °C for 30 seconds, followed by incubation at 72 °C for 2 minutes, and 4 °C hold. Samples were purified using the QIAquick PCR Purification Kit (Qiagen, 28104) and eluted in 30 μL EB. The length and purity of the samples were visualized on a 2% EX E-gel (ThermoFisher, G401002). A mutant for the ribozyme reaction template A8U was made by annealing 12 μL each of phi25_ag_R0_A8U and phi25_ag_R0_A8U_RC, each at 100 ng/μL, in 16 μL of 5X Reaction Buffer from the TranscriptAid T7 High Yield Transcription Kit (ThermoFisher, K0441). The sample was incubated at 94 °C for 2 minutes and cooled to room temperature before adding 32 μL NTP mix and 8 μL T7 RNA polymerase enzyme mix from the transcription kit. This sample was then *in vitro* transcribed following the same method as the following five samples.

Ribozyme constructs were *in vitro* transcribed following the TranscriptAid T7 High Yield Transcription Kit. Two reactions were made per sample, and 12 μL of purified DNA was used per construct. The reactions were incubated at 37 °C for 30 minutes, and DNase treated at 37 °C for 15 minutes. All samples were then purified following the RNA Clean & Concentrator-25 Kit (Zymo, R1017) and eluted in 30 μL water. All RNA samples were then purified on a denaturing 10% polyacrylamide gel made from 40% Acrylamide/Bis 29:1 solution (Bio-Rad, #1610146) in 1X TBE and 7 M urea, cast in an 18-well Criterion Empty Cassette (Bio-Rad, #3459902). RNA samples were mixed with 2X loading dye containing formamide and heated at 90 °C for 3 minutes before loading onto the 10% gel. The gel was run at 18 W for 45 minutes and stained with SYBR Gold (ThermoFisher, S11494). Samples were extracted from the gel following the ZR small-RNA PAGE Recovery Kit (Zymo, R1070) and eluted in 25 μL of water. Concentrations were measured using a NanoDrop and the length and purity was confirmed by the Agilent Bioanalyzer.

#### Polymerization reactions

Polymerization timecourses were carried out with final concentrations of 1 μM ribozyme, 0.5 μM template, 0.4 μM primer, 200 mM MgCl_2_, 4 mM each NTP, 50 mM Tris-HCl, pH 8.3, and 0.05% TWEEN 20 at 17 °C. Each extension reaction was prepared by mixing 2 μL 1 M Tris-HCl pH 8.3, 0.4 μL water, 10 μL 4 μM ribozyme, 10 μL 2 μM template RNA, and 1.6 μL 10 μM fluorescently labeled primer, chemically synthesized and purified with RNAse-free HPLC purification by IDT (**Table S2**). Template RNA R0 was synthesized by IDT. The reactions were heated to 80 °C and cooled to 17 °C over 5 minutes in the thermocycler. To each reaction, the following was added to give a final volume of 40 μL: 1.4 μL water, 0.2 μL 10% TWEEN 20, 6.4 μL 25 mM NTP mixture, 8 μL 1 M MgCl_2_. The reactions were incubated at 17 °C and 3 μL was taken out from each reaction at the time-points and quenched with 1.5 μL 0.5 M EDTA. The quenched samples were purified by ethanol precipitation by adding 1 μL GlycoBlue (ThermoFisher, AM9516), 20 μL 3 M sodium acetate pH 5.5, 174.5 μL water, mixed well before adding 600 μL 100% ethanol and incubated on dry ice for 30 minutes. The samples were centrifuged at 21,000×g for 30 minutes at 4 °C to pellet the sample. The supernatant was decanted and the pellet was washed with 600 μL cold 70% ethanol and centrifuged at 21,000×g for 3 minutes at 4 °C. The ethanol wash step was repeated one more time and the residual ethanol was removed by air drying the pellet for 10 minutes. Each pellet was resuspended in 10 μL water.

Samples from the time course were electrophoresed on a denaturing 20% polyacrylamide gel made from 20% Acrylamide-UREA Solution (IBI Scientific, IB70012) in 1X TBE and 7 M urea, cast in a 26-well Criterion Empty Cassette (Bio-Rad, #3459903). 2.5 μL sample was mixed with 7.5 μL loading dye containing formamide and heated at 90 °C for 3 minutes before loading onto the 20% gel. The gel was electrophoresed at 14 W for 45 minutes and imaged on the Typhoon FLA 9500 (Cytiva) under FAM or Cy5 fluorescence.

#### Analysis of polymerization kinetics

Densitometric measurements of time course gel scans were taken with the ImageQuant™ TL analysis software 11.0 (Cytiva Life Sciences). The percentage of unextended primer was calculated at each time point and the data were plotted in GraphPad Prism 11 (Graphpad Software, Inc.). The depletion of the unextended primer was used to estimate the rate of first nucleotide addition. In all cases, the data were fit by a one-phase association model:

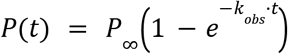

Where *p*(*t*) is the percentage of extended primer at time *t, p*_∞_ is the percentage of extended primer as *t* approaches infinity, and so *k*_obs_ is the observed rate of the addition of the first nucleotide to the unextended primer. Confidence intervals reflect fitting uncertainties. Our current measurements do not separate *k*_obs_ into microscopic rate constants and cannot be interpreted in terms of microscopic steps (ribozyme equilibration upon addition of MgCl_2_ and NTPs, template binding, docking, catalysis, extension at each template nucleotide, etc.); in particular, reported effects report on the extension of the first nucleotide and not later extension steps.

For double-mutant analyses, uncertainty propagation was carried out on ln *k*_obs_ rather than on *k*_obs_, since fitting errors in rate constants were approximately log-normally distributed. Ratios of rate constants were computed as differences of logarithms, with standard deviations combined in quadrature, confidence intervals recovered as ± 1.96σ, followed by exponentiation to return to *k*_obs_.

For each pair of mutations, the rate relative to WT if the two acted independently is (*k*_1_ / *k*_WT_) × (*k*_2_ / *k*_WT_) and the cooperativity factor Ω, equivalent to the nonadditivity term of a double-mutant cycle (*41, 42*), is

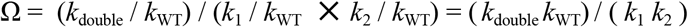

Ω = 1 indicates independent effects; Ω > 1 indicates positive cooperativity, the signature of an energetic interaction between the two mutated positions. Uncertainty in ln Ω was obtained by combining the standard deviations of all four measurements in quadrature. An apparent coupling free energy may be obtained as ΔΔG_int_ = −RT ln Ω, e.g., corresponding to −0.88 kcal mol^−1^ [95% CI: −0.99 to −0.77 kcal mol^−1^] for the C31:G162 cycle at 17 °C. Reaction of the A8U template with WT ribozyme gave weak activity, using either an *in vitro* transcribed template with a prepended AG (ag_R0_A8U; **Table S2**) or chemically synthesized R0_A8U; extension products were limited primarily to single-nucleotide extension and the resulting band overlapped with the unextended band, allowing only an upper bound on activity. To give the most conservative estimates of cooperativity, the main text and **Table 1** gives upper bounds for this reaction rate, derived from the highest values estimated (with chemically synthesized R0_A8U). Reaction of the A8U template with U2A ribozyme gave rapid reaction rates; to give the most conservative estimates for this increase, the main text and Table 1 give lower values (1.6-fold with in vitro transcribed ag_R0_A8U, compared to 2.8-fold with chemically synthesized R0_A8U).

### Structure motif search

Three tertiary motifs were isolated from the cryo-EM structure: the NTP site (G35–U37, A100–A102 and A148–A150), the α/α′ contact (twelve residues, C29–G33 and G158–C164), and a six-residue base stack (A34, A62, A145, C146, A151 and A163) drawn from five sequence-separated regions. Each was searched against the Protein Data Bank with ARTEM 2.0 (*31*), which superimposes local sets of nucleotides independently of backbone connectivity and of base identity, using the BGSU RNA 3D Hub non-redundant list (*43*), release 4.50 (2026-07-29) at a 3.0 Å resolution cutoff.

### 3D modeling

Three dimensional models of the complete tC19Z *apo* structure and *holo* structure with primer, template, and NTP bound were prepared with Rosetta’s FARFAR method with the presented cryo-EM structure and class I ligase structure (PDB 3R1L) used as templates for homology modeling (*44, 45*). Residues G147, A148, and residues involved in the ectopic pairing (1-7 and 103-109) were removed and rebuilt during *holo* modeling.

## Use of AI-assisted technologies

Anthropic’s Claude (Opus 5.0) was used for assistance with prose editing; all AI-generated text and analyses were reviewed, edited, and verified by authors before incorporation. The authors take full responsibility for the accuracy of all content.

## Author Contributions

D.B.H. designed, purified, and prepared cryo-EM grids for the scaffolded tC19Z construct. N.S., A.M., and Z.Y. collected, processed and analyzed cryo-EM data. J.H., D.B.H., B.R., and N.T. modeled cryo-EM coordinates. V.W. and R.D. designed and carried out polymerization kinetics experiments, with advice from DS; DS and RD analyzed kinetics experiments. D.S., J.H., and R.D. prepared the manuscript draft with input and revisions from all authors.

## Data Availability

The cryo-EM structure of the tC19Z RNA polymerase ribozyme in its apo state is available in the PDB under accession ID 37SO. Cryo-EM maps are available at the EMDB under accession ID EMDB-78467.

## Supplementary Materials

Supplementary PDF containing Supplementary Text, Tables S1 to S2, Figs. S1 to S5.

Movie S1: Global structure of tC19Z fit into map, followed by overlay with class I ligase.

Movie S2: Tertiary elements at the interface of the catalytic and accessory domains of tC19Z.

Movie S3: Magnesium ions form a seam at the interface of the catalytic and accessory domains. Movie S4: Conformational change required between apo and holo structures of tC19Z.

File S1: Model of complete apo tC19Z molecule using FARFAR2, in PDB format.

File S2: Model of complete holo tC19Z molecule with primer, template, and NTP using FARFAR2, in PDB format.

## Acknowledgements

We thank I. Zheludev, K. Zhang, and W. Chiu (Stanford) for preliminary efforts in structural and functional characterization of RNA polymerase ribozymes; E. Baulin (IMol) for advice on ARTEM motif search; G. Meissner (Janelia, HHMI) and AI@HHMI for support; and G. Joyce, D. Horning, and the Joyce lab (Salk) for discussions of the structure. We acknowledge funding from the National Institutes of Health (R35GM122579 to R.D.; R35GM141706 to N.T.), National Science Foundation (2330652 to R.D.), and the Howard Hughes Medical Institute (R.D.). Claude (Opus 5.0) was used for editorial assistance during manuscript preparation; all AI-assisted output was reviewed and verified by the authors.

