## Supplementary Text for "Cryo-EM of a nucleotide-polymerizing ribozyme enables its predictive improvement"

### **Supplementary Materials** for "Cryo-EM of a nucleotide-polymerizing ribozyme enables its predictive improvement"

This document contains the following sections:

**Supplementary Text**
Secondary structure of the accessory domain
Additional interactions in α-α′
Hypothesis for partial rescue in α-α′ compensatory mutagenesis
Structural precedents for tC19Z tertiary motifs
Correspondence between the proposed tC19Z NTP binding site and the class I ligase substrate binding site
**Supplementary Tables**
**Supplementary Figures**
**Supplementary References**

### Supplementary Text

#### Secondary structure of the accessory domain

While the catalytic domain of tC19Z is largely conserved from the ancestral class I ligase, the accessory domain represents the major structural addition associated with polymerase function. The secondary structure of the accessory domain matches previous predictions based on covariation and chemical mapping (*1*, *2*). This secondary structure, shared among Pol 1-derived accessory domains, can be described with two stems: A3 and A4 (using the nomenclature of Wang *et al.* 2011 (*2*)), interrupted by an A-rich loop (previously also called the 'purine-rich bulge' (*2*)). A4 also has a characteristic single-nucleotide bulge whose presence is conserved but whose identity is not conserved (**Figure 1C**). Although the A4 stem was originally predicted to be capped by a triloop (*3*), chemical mapping suggested it to be a pentaloop (*2*), which was confirmed here (**Figure 1C** and below). Beyond the A3 and A4 stems, the accessory domain secondary structure also includes two additional predicted helices (A1 and A2, **Figure 1C**), which were weakly resolved in the map and are dispensable for the activity of Pol 1-derived ribozymes, and so are not discussed further.

#### Additional interactions in α-α′

The α-α′ is reminiscent of a T-loop motif (*4*), this interaction begins with a non-canonical Watson-Crick:Hoogsteen U:A base pair at the base of the A4 pentaloop; however, unlike the conventional

T-loop motif, the adenosine residue in this interaction does not come from within the same loop but instead belongs to the triloop in the J3/4 region of the catalytic domain (**Figure 2B**).

The α-α′ interaction involves two peripheral base-stacking interactions (**Figure 2B**). Although the identity of the nucleotide at position 156 is not strictly required to be an adenosine, it is universally conserved as a single-nucleotide bulge (*1–3*, *5–9*). The structure reveals that this bulged nucleotide stacks with U30 (another nucleotide whose position but not identity is not conserved) on the opposite end of the J3/4 triloop, clamping the J3/4 triloop against the A4 stem. A second contact is made by A163 in the A4 pentaloop, which hydrogen bonds and stacks with A62 as part of the long-range hexastack (**Figure 2F**). Together, these interactions enclose the J3/4 triloop within A4 and its pentaloop, suggesting that the A4 stem of the accessory domain functions as an aptamer for the J3/4 region of the catalytic domain. This interpretation is supported by earlier work showing that the two domains can assemble *in trans* (*10*).

In addition, C31, A32, G33, G158, and U159 from α-α′ are all implicated in the hydrated magnesium seam. C31 and A32 each contribute one non-bridging phosphate oxygen to the inner-shell coordination of a $Mg^{2+}$ ion, while G33 appears to hold a central role in this region, with its non-bridging phosphate oxygens each coordinating an additional $Mg^{2+}$ ion. The nucleobases of G158 and U159 hydrogen bond to water molecules coordinated by the $Mg^{2+}$ ion held by C31 and A32 (**Figure 2E**).

### Hypothesis for partial rescue in α-α′ compensatory mutagenesis

Cooperativity in double mutant analyses inform on interactions of the mutated residues with each other; partial rescue is a signature that the residues have additional interactions with other residues not included in the double mutant analysis. On observing partial rescue in the double mutant analysis of C31:G162 pair, we noted that the α-α′ interaction include two potential hydrogen bonding interactions between the sugar and phosphate of U160 and the Hoogsteen edge of the C31:G162 pair (**Figure 2B**). Notably, all three of the tested mutants would have some, or all of these potential hydrogen bonds disrupted, which may explain the double mutant's inability to fully rescue activity (**Figure 2C, 2D**), indicating that this Watson-Crick pairing is only partially responsible for the stability of this tertiary interaction. If the hydrogen bond between the 2′-hydroxyl of U160 and the N7 of G162 are particularly important for stabilizing the α-α′ interaction, this could also explain why the lone C31G mutation had a modest effect (2-fold; **Table 1**) on polymerase activity.

### Structural precedents for tC19Z tertiary motifs

To ask whether the tertiary elements seen in the tC19Z cryo-EM structure have precedents among known RNA structures, we searched three motifs against a non-redundant set of 2,600 RNA structures using ARTEM (*11*), which detects local nucleotide arrangements without regard to backbone connectivity or base identity (**Methods**). Because ARTEM scores base geometry alone, we also annotated every match with the number of contiguous sequence runs it spans. This annotation distinguished genuine motif recurrence from the background of geometric coincidences that a purely spatial search returns.

*The α-α′ contact is novel.* The twelve-residue α-α′ motif, composed of α triloop and the α′ pentaloop with their closing base pairs returned no match within 2.0 Å RMSD, at either eleven or twelve matched residues. Relaxing the threshold to recover the nearest arrangements anywhere in the database yielded closest matches to the same residue collection in both the *N. crassa* mitoribosomal small subunit rRNA (PDB 6YW5, 2.299 Å) and the *V. natriegens* ribosomal small subunit rRNA (PDB 9H90, 2.305 Å) (**Fig. S5A**). Both satisfied the geometry with seven disconnected fragments, as compared to the motif's two strands. This interaction therefore appears to be without structural precedent in the non-redundant PDB.

*The hexastack is geometrically common but topologically unique.* The hexastack resembles a single strand of an A-form helix but is assembled from five sequence-separated segments in tC19Z. Searching at a flat 2.0 Å cutoff gave 697 matches, most of which came from a single contiguous segment; only three matched the tC19Z hexastack five-region topology: an 80S ribosome (PDB 8Q87, 1.778 Å), an RNP complex (PDB 5WTK, 1.912 Å), and a hairpin ribozyme (PDB 3BBM, 1.987 Å) (**Fig. S5B**). All three lay near the 2.0 Å RMSD cutoff and missed one stacking interaction. We therefore found no convincing precedent for the hexastack arrangement.

*The proposed NTP site matches the class I ligase ribozyme*. The nine-residue proposed NTP site (**Fig. 2G** and table below) returned 4,265 matches across 307 non-redundant classes within 2 Å. The single strongest match was PDB 3HHN, the representative in the non-redundant set of the class I ligase ribozyme, which is the ancestor of the tC19Z polymerase lineage. Eight of the nine motif residues superimposed at 0.563 Å RMSD, and the correspondence preserved the motif's architecture of three sequence-separated strands (**Fig. S5C** and see next section). The next closest match (4L81, a SAM I/IV riboswitch) displayed a similar A-minor interaction (**Figure S5C**), but did not match the C36 bulge and gave a higher RMSD (1.077 Å) over the remaining nucleotides.

#### Correspondence between the proposed tC19Z NTP binding site and the class I ligase substrate binding site

The proposed tC19Z NTP binding site and the substrate binding site of the class I ligase are constructed from residues at different positions in the two molecules, and from different domains. Eight of the nine residues comprising the tC19Z site superimpose on the PDB representing the cIL in our non-rednudant set (PDB 3HHN, chain E) at 0.56 Å RMSD with base identity conserved at every matched position (**Methods**, **Supplementary Text: Structural precedents for tC19Z tertiary motifs**). Correspondences are given below; "cIL-" denotes class I ligase numbering throughout.

| tC19Z | tC19Z domain | cIL (3HHN) | Interaction |
|---|---|---|---|
| G35 | catalytic | cIL-G46 | A-minor acceptor pair with C101 / cIL-C113 |
| C36 | catalytic | cIL-C47 | — |
| U37 | catalytic | cIL-U48 | — |
| A100 | catalytic | cIL-A112 | — |
| C101 | catalytic | cIL-C113 | A-minor acceptor pair with G35 / cIL-G46 |

| | | | |
|---|---|---|---|
| A102 | catalytic | cIL-A114 | Stacking with A150 / cIL-A4 |
| A148 | accessory | (cIL-G2) | Backbone superposition only; base not matched |
| A149 | accessory | cIL-A3 | A-minor donor to G35:C101 / cIL-G46:cIL-C113 |
| A150 | accessory | cIL-A4 | Splayed backbone with A149; stacking on A102 / cIL-A114 |

The three residues that follow the ligase's 5′ triphosphorylated nucleotide (cIL-G1) in the ligation reaction (cIL-G2, cIL-A3, and cIL-A4) were excised during the selections that produced the polymerase lineage. The functions of cIL-A3 and cIL-A4 are recovered in tC19Z by A149 and A150, which reach the same site from the opposite end of the molecule. A148 recovers the backbone position of cIL-G2 but not its base orientation, and was not matched in the ARTEM superposition.

Beyond the correspondences above, modeling an NTP into the tC19Z site by superposition with the substrate-bound ligase placed it in slight steric conflict with G147, the residue immediately preceding the A148–A150 segment and without correspondence to nucleotides in the class I ligase. G147 is therefore expected to occupy a different position in the substrate-bound state, although this map does not indicate where. Consistent with an intrinsically mobile residue, density for G147 is weak in our map.

### Supplementary Tables

| | **tC19Z apo**<br>(PDB ID: 37SO, EMD-78467) |
|---|---|
| **Data collection and Processing** | |
| Microscope | Titan Krios G4 |
| Voltage (keV) | 300 |
| Camera | Falcon 4i |
| Magnification | 165,000x |
| Pixel size at detector (Å/pixel) | 0.7336 |
| Total electron exposure ($e^-$/Å$^2$) | 50 |
| Exposure rate ($e^-$/pixel/sec) | 8.45 |
| Number of frames collected during exposure | 1017 |
| Defocus range (μm) | -0.4 to -2.0 |
| Automation software (EPU, SerialEM or manual) | SerialEM |
| Energy filter slit width | 6 eV |
| Micrographs collected (no.) | 10,892 |
| Micrographs used (no.) | 10,443 |
| Total extracted particles (no.) | 1,094,542 |
| Refined particles (no.) | 152,136 |
| Final particles (no.) | 150,593 |
| Point-group or helical symmetry parameters | C1 |
| Resolution (global, Å) | |
| FSC 0.5 (unmasked/masked) | 6.4/3.9 |
| FSC 0.143 (unmasked/masked) | 4.3/3.1 |
| Resolution range (local, Å) | 2.64 - 13.33 |
| Resolution range due to anisotropy (Å) | 2.96 - 3.73 |
| Map sharpening *B* factor (Å$^2$) / (B factor Range) | 85.4 |
| Map sharpening methods | Global |
| **Model composition** | |
| RNA (nt) | 165 |
| $Mg^{2+}$ | 5 |
| Water | 25 |
| **Model Refinement** | |
| Refinement package | PHENIX |
| - real or reciprocal space | Real |
| - resolution cutoff | 3.1 |
| Model-Map scores | |
| - CC | 0.85 |
| - Average FSC | 0.72 |
| *B* factors (Å$^2$) | |
| RNA | 79.06 |
| $Mg^{2+}$ | 9.51 |
| Water | 9.01 |
| R.m.s. deviations from ideal values | |
| Bond lengths (Å) | 0.007 |

| | |
|---|---|
| Bond angles (°) | 0.716 |
| **Validation** | |
| MolProbity score | 2.44 |
| Clashscore | 5.45 |

**Table S1.** Cryo-EM data collection, refinement, and validation statistics

| Name | Sequence |
| --- | --- |
| **Cryo-electron microscopy** | |
| tC19Z cryo-EM construct DNA, including group II intron scaffold and BamHI | TAATACGACTCACTATAGTCATTGAAAAAAAAAGACAAATCTGCCCTCAGAGCTT<br>GAGAACATC*CAGAAGTCAGCAGAAGTCATAGTACCCTTCGGGGGAAGGACGGAACA*<br>*AGTATGGCGTTCGCGCCATGCTTGAACCACCGTATACCGAACGGTACGTACGGTGGTG*<br>*AAACAAACAAATAAACTAAATTATGTGTGCCCGGCATGGGTGCAGTCTATAGGGTGAGA*<br>*GTCCCGAACTGTGAAGGCAGAAGTAACAGTTAGCCTAACGCAAGGGTGTCCGTGGCG*<br>*ACATGGAATCTGAAGGAAGCGGACGGCAAACCTTCGGTCTGAGGAACACGAACTTCA*<br>*TATGAGGCTAGGTATCAATGGATGAGTTTGCATAACAAAACAAAGTCCTTTCTGCCAAA*<br>*GTTGGTACAGAGTAAATGAAGCAGATTGATGAAGGGAAAGACTGCATTCTTACCCGGG*<br>*GAGGTCTG*GATGCAGAGGAGGCAGCCTTCGGTGGCGCGATAGCGCCAACGTTCT<br>CAACAGACACCCAATACTCCCGCTTCGGCGGGTGGGGATAACACCTGACGAAAA<br>GGCGATGTTAGACACGCCCAGGTCATAATCCCCGGAGCTTCGGCTCC*GGATCC* |
| tC19Z cryo-EM construct RNA | GUCAUUGAAAAAAAAAGACAAAUCUGCCCUCAGAGCUUGAGAACAUC*CAGAA*<br>*GU*CAGC*AGAAGUCAUAGUACCCUUCGGGGGAAGGACGGAACAAGUAUGGCGUUC*<br>*GCGCCAUGCUUGAACCACCGUAUACCGAACGGUACGUACGGUGGUGAAACAAACA*<br>*AAUAAACUAAAUUAUGUGUGCCCGGCAUGGGUGCAGUCUAUAGGGUGAGAGUCC*<br>*CGAACUGUGAAGGCAGAAGUAACAGUUAGCCUAACGCAAGGGUGUCCGUGGCGA*<br>*CAUGGAAUCUGAAGGAAGCGGACGGCAAACCUUCGGUCUGAGGAACACGAACUUC*<br>*AUAUGAGGCUAGGUAUCAAUGGAUGAGUUUGCAUAACAAAACAAAGUCCUUUCUG*<br>*CCAAAGUUGGUACAGAGUAAAUGAAGCAGAUUGAUGAAGGGAAAGACUGCAUUCU*<br>*UACCCGGGGAGGUCUG*GAUGCAGAGGAGGCAGCCUUCGGUGGCGCGAUAGCG<br>CCAACGUUCUCAACAGACACCCAAUACUCCCGCUUCGGCGGGUGGGGAUAAC<br>ACCUGACGAAAAGGCGAUGUUAGACACGCCCAGGUCAUAAUCCCCGGAGCUU<br>CGGCUCC*GGAUC* |
| **Polymerization reactions** | |
| tC19Z_wt DNA | TTCTAATACGACTCACTATTAGTCATTGAAAAAAAAAGACAAATCTGCCCTCAGA<br>GCTTGAGAACATCTTCGGATGCAGAGGAGGCAGCCTTCGGTGGCGCGATAGCGC<br>CAACGTTCTCAACAGACACCCAATACTCCCGCTTCGGCGGGTGGGGATAACACC<br>TGACGAAAAGGCGATGTTAGACACGCCCAGGTCATAATCCCCGGAGCTTCGGCT<br>CC |
| tC19Z_G162C DNA | TTCTAATACGACTCACTATTAGTCATTGAAAAAAAAAGACAAATCTGCCCTCAGA<br>GCTTGAGAACATCTTCGGATGCAGAGGAGGCAGCCTTCGGTGGCGCGATAGCGC<br>CAACGTTCTCAACAGACACCCAATACTCCCGCTTCGGCGGGTGGGGATAACACC<br>TGACGAAAAGGCGATGTTACACACGCCCAGGTCATAATCCCCGGAGCTTCGGCT<br>CC |
| tC19Z_C31G DNA | TTCTAATACGACTCACTATTAGTCATTGAAAAAAAAAGACAAATCTGCCCTGAGA<br>GCTTGAGAACATCTTCGGATGCAGAGGAGGCAGCCTTCGGTGGCGCGATAGCGC<br>CAACGTTCTCAACAGACACCCAATACTCCCGCTTCGGCGGGTGGGGATAACACC<br>TGACGAAAAGGCGATGTTAGACACGCCCAGGTCATAATCCCCGGAGCTTCGGCT<br>CC |
| tC19Z_C31G_G162C DNA | TTCTAATACGACTCACTATTAGTCATTGAAAAAAAAAGACAAATCTGCCCTGAGA<br>GCTTGAGAACATCTTCGGATGCAGAGGAGGCAGCCTTCGGTGGCGCGATAGCGC<br>CAACGTTCTCAACAGACACCCAATACTCCCGCTTCGGCGGGTGGGGATAACACC<br>TGACGAAAAGGCGATGTTACACACGCCCAGGTCATAATCCCCGGAGCTTCGGCT<br>CC |
| tC19Z_U2A DNA | TTCTAATACGACTCACTATTAGACATTGAAAAAAAAAGACAAATCTGCCCTCAGA<br>GCTTGAGAACATCTTCGGATGCAGAGGAGGCAGCCTTCGGTGGCGCGATAGCGC |

| | |
|---|---|
| | CAACGTTCTCAACAGACACCCAATACTCCCGCTTCGGCGGGTGGGGATAACACCTGACGAAAAGGCGATGTTAGACACGCCCAGGTCATAATCCCCGGAGCTTCGGCTCC |
| phi25_ag_R0_A8U | TTCTAATACGACTCACTATTAGGACAATGTCAAAAAACACTCACACACACTCCACACCCTCGTTTTGCTCCGAC |
| phi25_ag_R0_A8U_RC | mGmUCGGAGCAAAACGAGGGTGTGGAGTGTGTGTGAGTGTTTTTTGACATTGTCCTAATAGTGAGTCGTATTAGAA |
| phi25_a_tC19Z-FIX-fwd | TTCTAATACGACTCACTATTAGTCATTG |
| phi25_a_tC19Z-FIX-fwd_U2A | TTCTAATACGACTCACTATTAGACATTG |
| tC19Z-rev_methyl | mGmGAGCCGAAGCTCCGGGGAT |
| tC19Z_wt RNA | AGUCAUUGAAAAAAAAAGACAAAUCUGCCCUCAGAGCUUGAGAACAUCUUCGGAUGCAGAGGAGGCAGCCUUCGGUGGCGCGAUAGCGCCAACGUUCUCAACAGACACCCAAUACUCCCGCUUCGGCGGGUGGGGAUAACACCUGACGAAAAGGCGAUGUUAGACACGCCCAGGUCAUAAUCCCCGGAGCUUCGGCUCC |
| tC19Z_G163C RNA | AGUCAUUGAAAAAAAAAGACAAAUCUGCCCUCAGAGCUUGAGAACAUCUUCGGAUGCAGAGGAGGCAGCCUUCGGUGGCGCGAUAGCGCCAACGUUCUCAACAGACACCCAAUACUCCCGCUUCGGCGGGUGGGGAUAACACCUGACGAAAAGGCGAUGUUACACACGCCCAGGUCAUAAUCCCCGGAGCUUCGGCUCC |
| tC19Z_C32G RNA | AGUCAUUGAAAAAAAAAGACAAAUCUGCCCUGAGAGCUUGAGAACAUCUUCGGAUGCAGAGGAGGCAGCCUUCGGUGGCGCGAUAGCGCCAACGUUCUCAACAGACACCCAAUACUCCCGCUUCGGCGGGUGGGGAUAACACCUGACGAAAAGGCGAUGUUAGACACGCCCAGGUCAUAAUCCCCGGAGCUUCGGCUCC |
| tC19Z_C32G_G163C RNA | AGUCAUUGAAAAAAAAAGACAAAUCUGCCCUGAGAGCUUGAGAACAUCUUCGGAUGCAGAGGAGGCAGCCUUCGGUGGCGCGAUAGCGCCAACGUUCUCAACAGACACCCAAUACUCCCGCUUCGGCGGGUGGGGAUAACACCUGACGAAAAGGCGAUGUUACACACGCCCAGGUCAUAAUCCCCGGAGCUUCGGCUCC |
| tC19Z_U2A RNA | AGACAUUGAAAAAAAAAGACAAAUCUGCCCUCAGAGCUUGAGAACAUCUUCGGAUGCAGAGGAGGCAGCCUUCGGUGGCGCGAUAGCGCCAACGUUCUCAACAGACACCCAAUACUCCCGCUUCGGCGGGUGGGGAUAACACCUGACGAAAAGGCGAUGUUAGACACGCCCAGGUCAUAAUCCCCGGAGCUUCGGCUCC |
| R0 RNA (IDT) | GACAAUGACAAAAAACACUCACACACACUCCACACCCUCGUUUUGCUCCGAC |
| R0_A8U RNA | AGGACAAUGUCAAAAAACACUCACACACACUCCACACCCUCGUUUUGCUCCGAC |
| R0_A8U RNA (IDT) | GACAAUGUCAAAAAACACUCACACACACUCCACACCCUCGUUUUGCUCCGAC |
| FAM-P1 | 5´-FAM-GGAGCAAAACGAGG |
| CY5-P1 | 5´-Cy5-GGAGCAAAACGAGG |

**Table S2.** DNA and RNA sequences used in this study. Underlined sequences indicate promoters for transcription by T7 RNA polymerase; italicized sequences indicate group II intron insert and BamHI restriction site in cryo-EM construct.

### Supplementary Figures

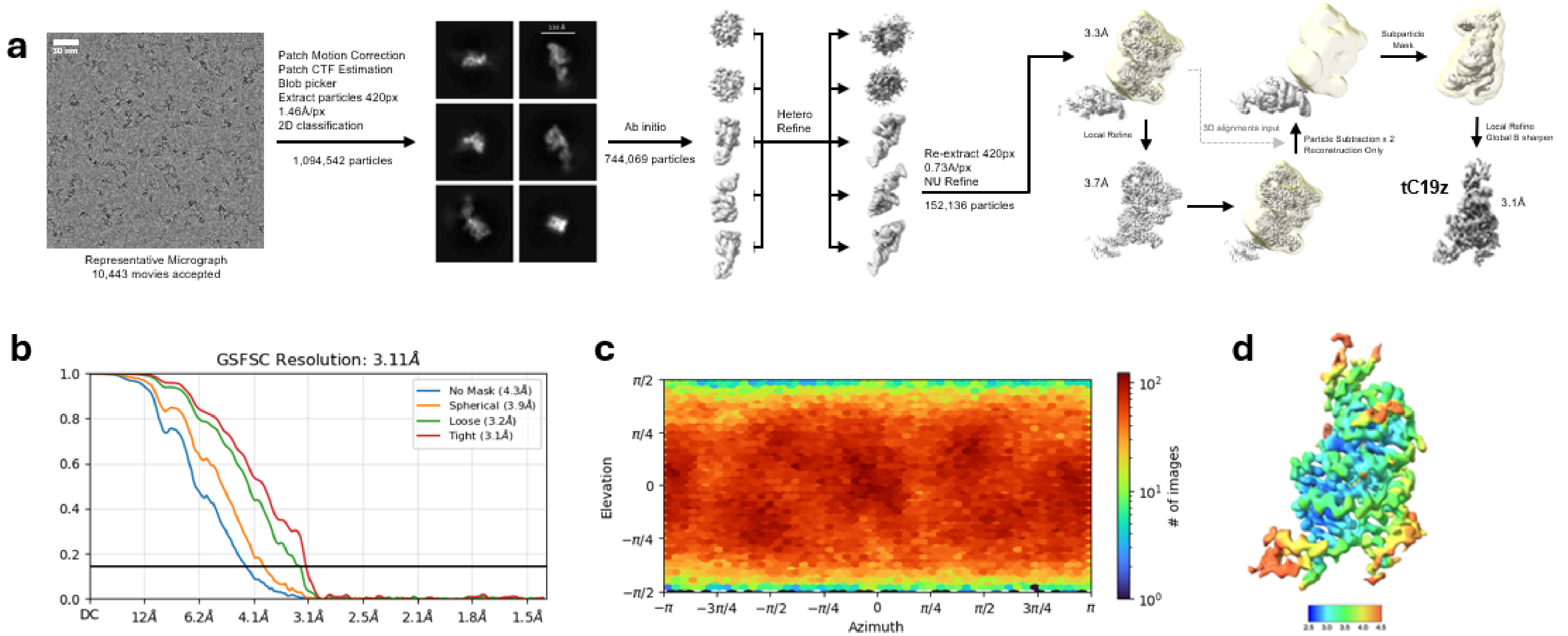

**Figure S1.** Cryo-EM data processing of tC19Z. **A)** Data processing workflow. **B)** Gold Standard Fourier Shell Correlation curve. **C)** Viewing direction distribution plot. **D)** Local resolution map.

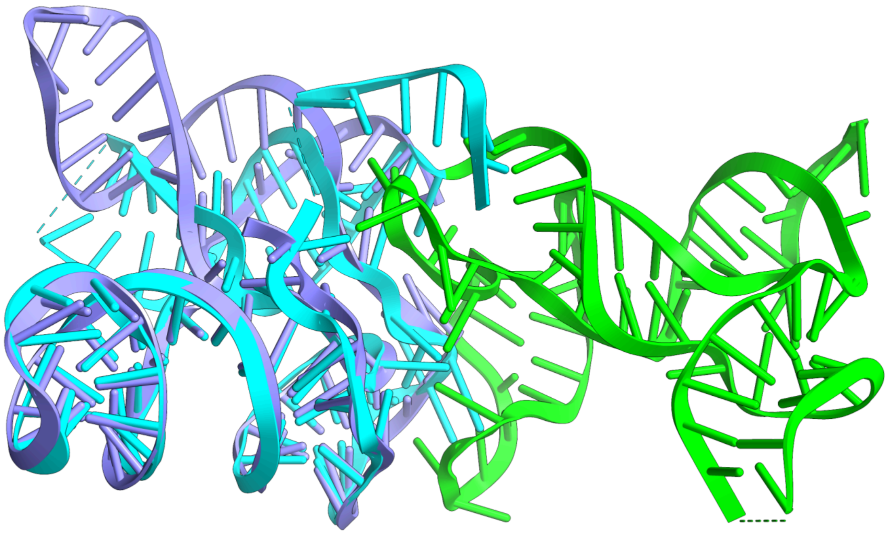

**Figure S2.** Overlay of structures of tC19Z (accessory domain in green, catalytic domain in cyan) and the catalytic 5TU subunit of the triplet polymerase ribozyme (PDB 8T2P, light purple) (*12*), showing retention of the catalytic core structure inherited from the class I ligase. Domains outside the catalytic domain of 5TU, including a separate non-catalytic t1 subunit that completes the 5TU heterodimer, are not shown.

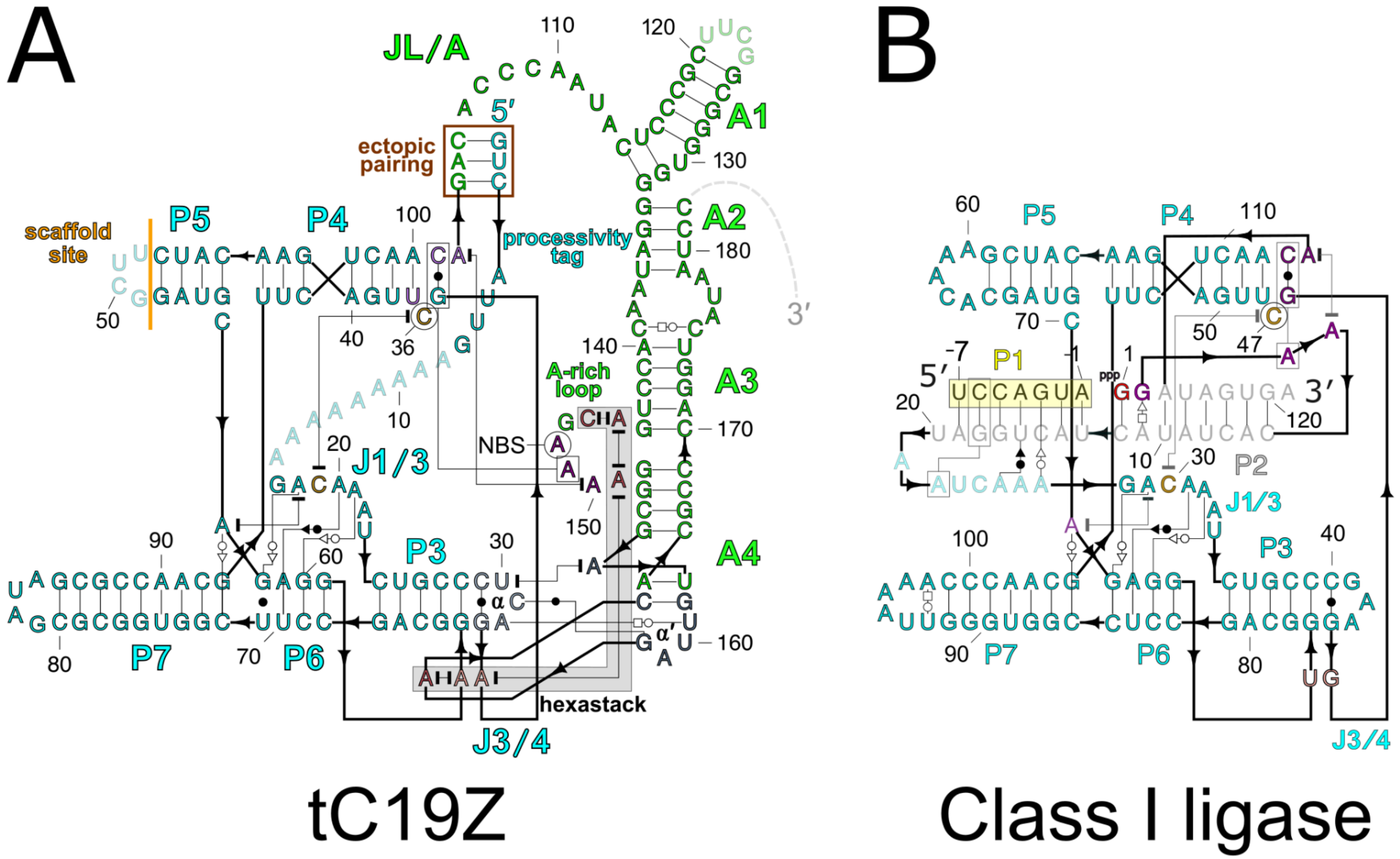

**Figure S3.** Secondary structure comparison of *apo* tC19Z and the class I ligase. A) Secondary structure of tC19Z, with colors as in **Figure 2**. B) Secondary structure of the class I ligase, adapted from PDB IDs 3HHN (*13*) and 3R1L (*14*). Analogs of gray nucleotides and primer (boxed in yellow) in the cIL structure are not visible in the *apo* tC19Z structure.

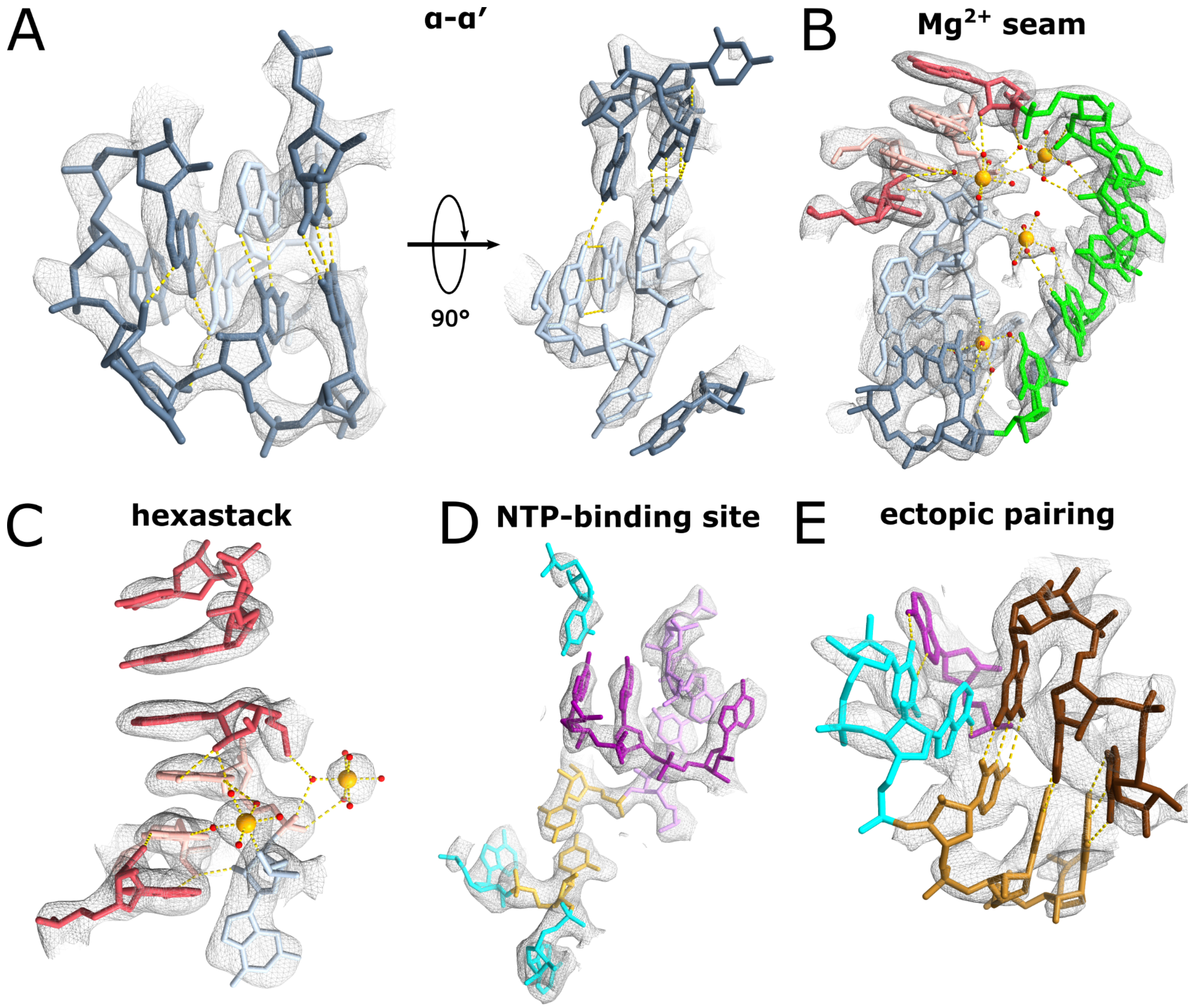

**Figure S4.** Close-up views of the interactions as seen in **Figure 2**, with cryo-EM map displayed. **A**) The α-α′ interaction as seen in **Figure 2B**, with the primary map shown at a level of 0.22. **B)** The $Mg^{2+}$ ion seam as seen in **Figure 2E**, with the alternate map shown at a level of 0.10 (supplementary map in EMDB-78467; this map has more detail in this region but lower resolution at the periphery). **C)** The hexastack as seen in **Figure 2F**, with the primary map shown at a level of 0.22. **D)** The proposed NTP binding site and active site as seen in **Figure 2H**, with the primary map shown at a level of 0.18. **E)** The ectopic pairing as shown in **Figure 2J**, with the primary map shown at a level of 0.15.

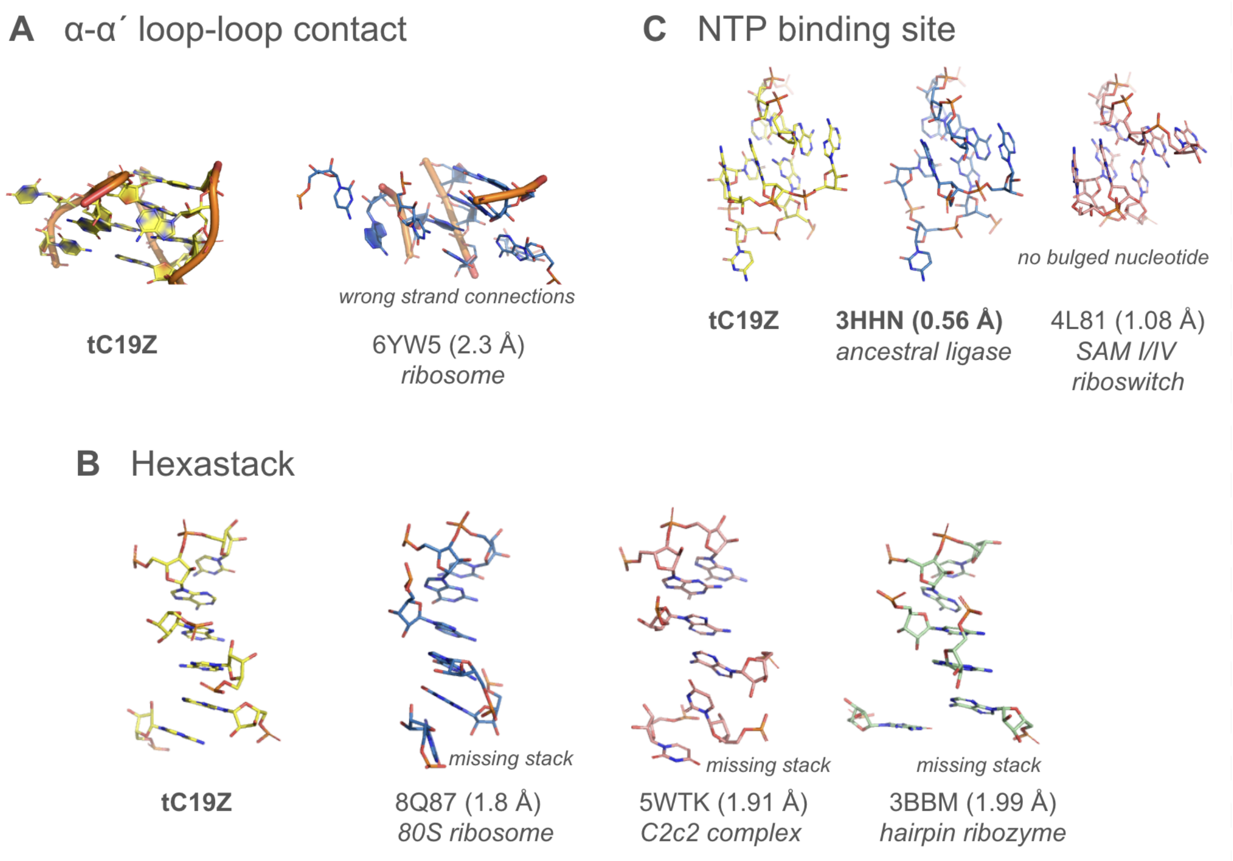

**Figure S5.** Comparison of closest structural matches in previously determined RNA structures to tertiary elements formed at the interface of the tC19Z catalytic and accessory domains: (A) α-α′ loop-loop contact, (B) hexastack, and (C) NTP binding site. Only the substrate binding site of tC19Z's ancestor, the class I ligase in (C) matches the strand connectivity and stacking pattern of any of these tC19Z elements.
